# Social interactions impact on the dopaminergic system and drive individuality

**DOI:** 10.1101/236554

**Authors:** N. Torquet, F. Marti, C. Campart, S. Tolu, C. Nguyen, V. Oberto, J. Naudé, S. Didienne, S. Jezequel, L. Le Gouestre, N. Debray, A. Mourot, J. Mariani, P. Faure

## Abstract

Individuality is a ubiquitous and well-conserved feature among animal species. The behavioral patterns of individual animals affect their respective role in the ecosystem and their prospects for survival. Even though some of the factors shaping individuality have been identified, the mechanisms underlying individuation are poorly understood and are generally considered to be genetics-based. Here we devised a large environment where mice live continuously, and observed that individuality, measured by both social and individual traits, emerged and settled within the group. Midbrain dopamine neurons underwent neurophysiological adaptations that mirrored this phenotypic divergence in individual behaviors. Strikingly, modifying the social environment resulted in a fast re-adaptation of both the animal’s personality and its dopaminergic signature. These results indicate that individuality can rapidly evolve upon social challenges, and does not just depend on the genetic or epigenetic initial status of the animal.

## Introduction

Individuality, or personality, refers to differences that remain stable over time and contexts for a series of behavioral traits expressed among individuals of the same species (Bach, 2009; Bergmüller and Taborsky, 2010; Duckworth, 2010; Sih et al., 2004; Wolf and Weissing, 2010). Individuality is a ubiquitous feature of animal populations (Pennisi, 2016). Evidence for phenotypic variability lead to extensive research on its adaptive significance and its ecological or evolutionary consequences (Dall et al., 2012; Gosling, 2001; Réale et al., 2007; Sih et al., 2004; van Overveld et al., 2013; Wolf and Weissing, 2010). Even though the proximal mechanisms underlying phenotypic variability could provide important information on the mechanisms underlying animal choices, stress responses or suseptibilities to disease, they have been understudied (Duckworth, 2010).

The emergence of animal personality has been linked to genetic and environmental interactions (Lynch and Kemp, 2014; Pennisi, 2016). Experiments with groups of near-clonal mice reared in a large and controlled environment have demonstrated behavioral divergence (Freund et al., 2013; Hager et al., 2014), which may emerge from the magnification of small initial differences in the epigenetic status or micro-environment of the animal (Lynch and Kemp, 2014). In this perspective, the combination of individual history and initial differences would form a unique path for each individual, and may explain the phenotypic variability observed at the population level. Social relationships are another factor with potential important roles in personality shaping. Notably, social stress studies identified susceptible and resilient animals (Berton et al., 2006; Krishnan et al., 2007), while social hierarchy analyses revealed that dominant animals are seemingly less sensitive to the effects of drugs than subordinates (Morgan et al., 2002). Normal or pathological social relationships can thus greatly modify individual behaviors in mice. However, the role of social relationships in the emergence of phenotypic variability is poorly understood. Interactions within a group were proposed to result in social specialization (Bergmüller and Taborsky, 2010), but whether the composition of a social group can affect non-social behavioral traits and the underlying neuronal processes remain to be determined.

Here we questioned the role of social relationships in the emergence of personality. For that purpose, we developed an experimental setup that combines an environment where animals live together, with a modular testing platform where animals are tested individually. In this environment, mice have individual access to specific feeding-related tasks while their social, circadian and cognitive behaviors are monitored continuously and for long periods of time using multiple sensors. This setup enabled the translation of activity and cognitive assessments into a definition of personality, and allowed to confirm the emergence of individuals with stable behavioral differences within a group of mice. Furthermore, we could demonstrate that individual’s traits correlate with neuronal activity at the level of the decision-making dopamine (DA) system. Finally, manipulating the social network was sufficient to reset both the animal’s personality and the activity of its DA cells. Altogether these data indicate that, in isogenic mice and for a conserved environment, social relationships govern individuality, most likely by impacting on the DAergic system.

## Results

### Automatic analysis of behavior in a large and naturalistic environment

Social life in natural environments and its consequences on the development of personality cannot be easily addressed in standardized behavioral laboratory tests. The development of automatic behavior analysis opens up new opportunities for in-depth phenotyping (Castelhano-Carlos et al., 2014; Endo et al., 2011; Krackow et al., 2010) and for studying individuation in the laboratory (Freund et al., 2013). An essential benefit of automation is the ability to conduct experiments on time scales that are orders of magnitude longer than traditional experiments (from minutes in classical assays to months of observation in automated systems). To test whether the social environment modifies individual traits, we first developed a complex and automatized environment, called “Souris City”, where male mice live in a group (10 to 20) for extended periods of time (2-3 months) while performing cognitive tests. Souris City is composed of a large environment (Social cage) connected to a test-zone where individual animals, isolated from their conspecifics, performed a test (here a choice task in a T-maze to obtain water, Fig. 1A and Supplementary Fig. 1). Animals were RFID-tagged and detected by antennas. This led to a coherent representation of mouse trajectories and distribution within the different compartments of Souris City: the nest compartment (NC), food compartment (FC), central compartment (CC), stairs (St) and T-maze (Fig. 1A). The circadian rhythm of the group emerged from pooled (n=49 mice; 5 experiments) activity measurement (Fig. 1B left). The time spent by mice in a given compartment generally varied from 1 to 30 min (Fig. 1B Right), with the shortest visits in FC, corresponding to feeding episodes. Conversely, very long stays (several hours) were found in NC, especially during the light time (Fig. 1C, see also Supplementary Fig. 2) and were associated with sleep episodes. These parameters described the general activity of the animals and can be used to construct more complex representations, such as the entropy of their distribution (see methods). Parameters describing group behavior can also be extracted, mainly using indicators that translate the simultaneous presence of a group of animals in a given space. As an example, a high rate of successive distinct RFID detections on a single antenna within short time intervals (<10 s) were indicative of social events, i.e. a group of mice passing from one compartment to another (Fig. 1D Left). Consecutive detections along different antennas (Fig. 1D Middle) indicated a follower tracking -or chasing- a leader (Fig. 1D right).

**Fig. 1:**
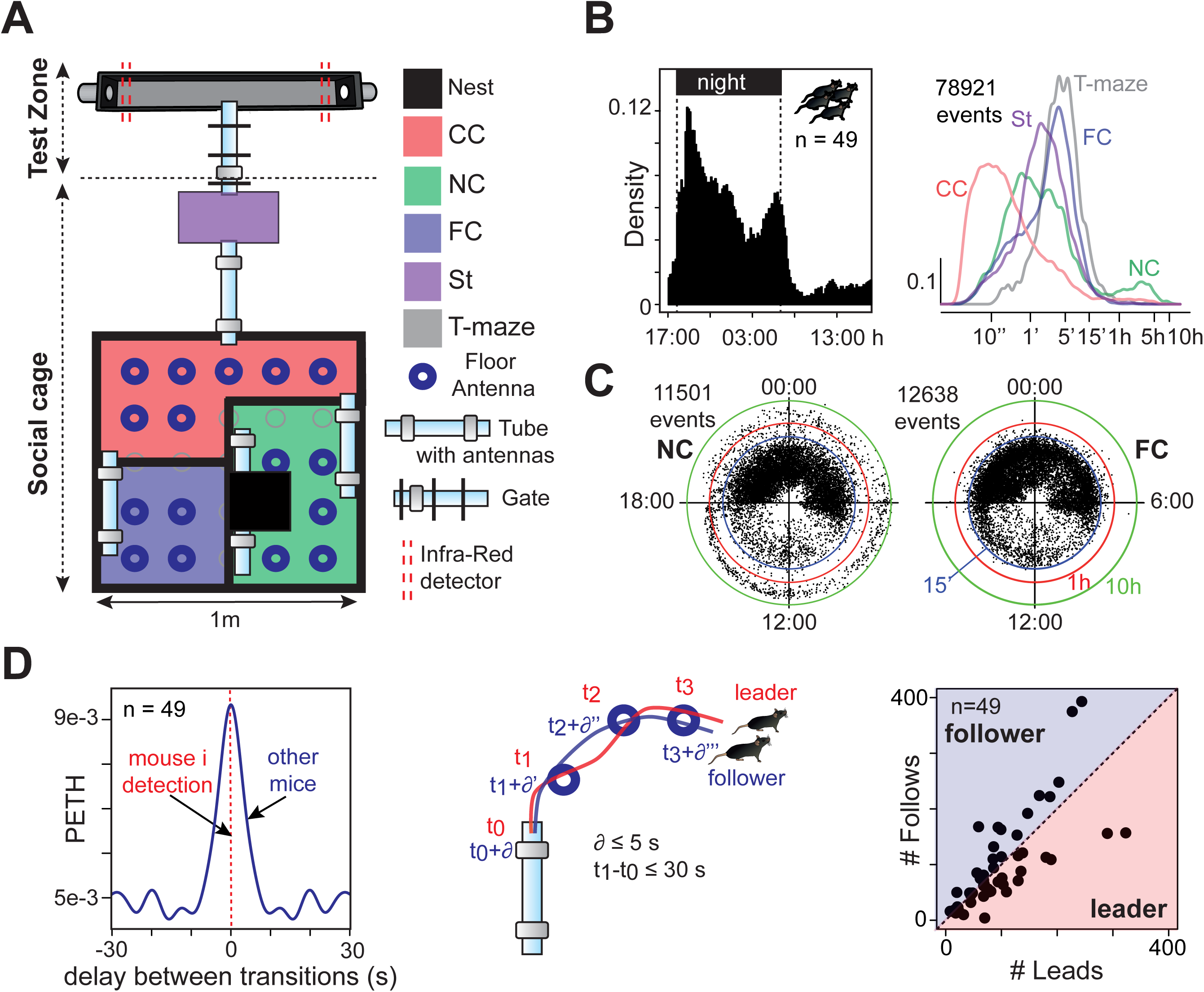
The Souris City environment. (**A**) Souris City setup with connectable compartments, gates and antennas. The setup is divided in two main parts: a social cage and a test zone. The social cage is divided in four compartments: NC, which contains a nest, FC where mice have free and uncontrolled access to food, CC and a stair (St) to get access to the gate (Supplementary Fig. 1). NC, FC and CC are located in a 1m x 1m square, on which St is connected by a tube. Mice are tagged with RFID chips and detected by floor or tubes RFID antennas. A gate separates the test zone (here a T-maze) from the social cage. Two infra-red beams (red dashed line) are used to detect mice in the T-maze. (**B**) (Left) Histogram of all the detection events from tubes (10 min time bins). (Right) Distribution of the time spent in each compartment (log-scale, bandwidth=0.1). (**C**) Circular plots showing the starting time (on a 24h dial) and duration (log distance of the point to the center) of each visit (a dot) for NC and FC. Three circles indicate the 15’ (blue), 1h (red) and 10h (green) limits. (**D**) Analysis of social behavior: (Left) Peri-event time histogram (PETH density, bandwidth = 2s) of detections from distinct mice on the same tube antenna, showing a delay between transitions lower than 10s. (Middle) Chasing episodes are defined by concomitant detections of the same two mice on at least two consecutive antennas. (Right) Follower and leader mice, based on the ratio between the number of leads over the number of follows. n=49 mice from 5 experiments.

### Evidence for the emergence of individual profiles in Souris City

Long-term exposition to complex and large social environments was shown to elicit a magnification of individual differences in groups of genetically identical mice (Freund et al., 2013). In agreement with this previous report, we observed in Souris City i) a large variety of behaviors, including atypical ones (Fig. 2A-B), and ii) the progressive divergence of individual measures linked to space occupancy, such as the entropy of animal distribution (Fig. 2B left), the time spent in a given compartment (Fig. 2B right) or the time spent alone (Supplementary Fig. 3A). These observations suggest a marked consistency in individual behaviors over time, which defines personality. To further substantiate the emergence of individuality, we quantified behavioral correlations upon context variations. We performed five sessions (Fig. 2C left) in which both the rules to access drink dispensers and the drinking solutions were modified. Indeed, access to the T-maze, and thus to the drink dispensers, can be controlled by a gate allowing the selective entry of one mouse at a time (see Supplementary Fig. 1). During the habituation period (Ha), mice explored Souris City and had free access to water (gate always open). Then, access was gate-restricted and the reward associated with drink delivery was modified along four sessions: water on both sides (session S1), water or sucrose 5% (S2), water or nothing (S3) and finally back to water on both sides (S4). Overall, such manipulations altered the territorial organization in the social cage with variations of space occupancy in the nest and stair compartments throughout the different sessions (Fig. 2C Middle and Right, Supplementary Fig. 3B). The modification of average behaviors across contexts contrasted with the stability of individual behaviors. For instance, animals spending less time than their conspecifics in the stair compartment in S1 correspondingly spent less time in this compartment in S2 (Fig. 2D), establishing a behavioral consistency throughout the experiment for any given animal. Similarly, a large set of behaviors showed strong homogeneity throughout the sessions, such as the animal inclination to lead or follow in chasing episodes (Fig. 2E), the proportion of time spent alone (Fig. 2F), or additional social and non-social individual traits (Fig. 2G, Supplementary Fig. 3C). Overall, our results establish that mice developed individual profiles in this large environment, i.e. they maintained unique and coherent behavioral trajectories throughout time and situations.

**Fig. 2:**
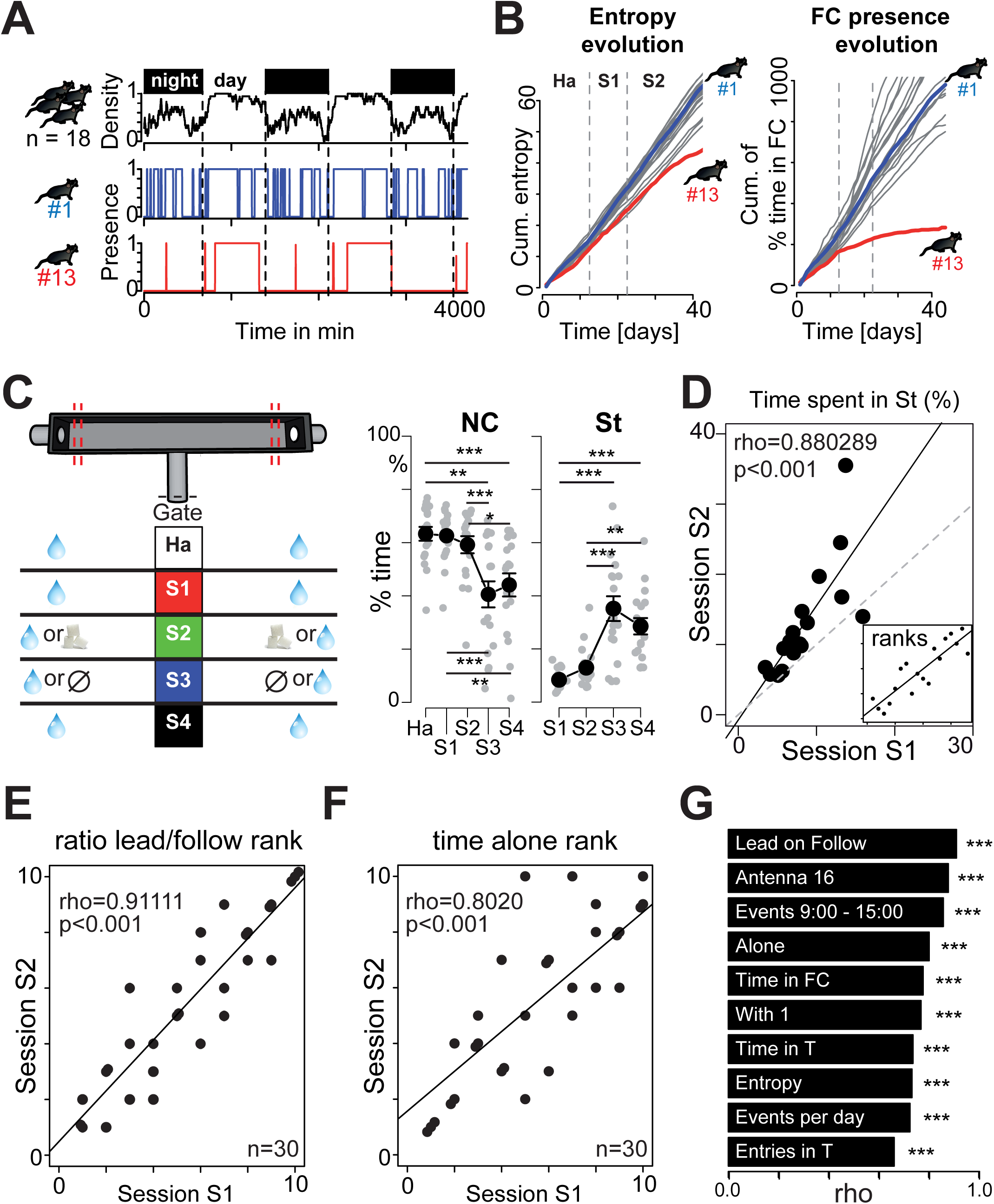
Consistency of behavior across situations. (**A**) Example of atypical behaviors. Top: density of mice in NC. Below: Presence (=1) of two mice (#1 and #13) in NC. (**B**) Cumulative distributions of entropy (Left) and of the proportion of time spent in FC (Right). (**C**) (Left) Diagram representing the five sessions. Variation across sessions (mean±sem) of (Middle) the proportion of time spent in NC (n=18, F(4,68)=22.69, p<0.001; and post-hoc test) and (Right) in St (n=18, F(3,51)=30.52, p<0.001; and post-hoc test). (D) Correlation between proportion of time spent in St for individual mice in session 1 (S1) against session 2 (S2) (Spearman correlation coefficient, n=18, fitted line= solid line, identity line= dotted line). Inset displays ranks instead of values with the correlation line. (**E-F**) Same as (D) inset for (**E**) the rank based on the ratio of leading over following and (**F**) the rank based on the proportion of time spent alone. (**G**). Rank correlations (rho) for two consecutive periods, for ten individual and social behaviors. For (e-f-g), n=30 mice from 3 independent experiments. ***p<0.001,**p<0.01,*p<0.05.

### Different strategies of decision-making outside the group

To refine individual description, we next addressed the relationship between social and nonsocial aspects of decision-making processes. In the T-maze with restricted access, mice voluntarily and individually performed a decision task, i.e. whether to make a left or right turn for accessing liquid reward. Once the choice for a particular arm (left or right) was made, the other arm closed off and the animal had to exit the test area for a new trial to begin (Supplementary Fig. 1D). The location of the different bottles was regularly swapped (every 3-4 days). The animal had thus to continually probe the environment and to adjust its behavior in response to changes in rewarding outcomes. The occupancy rate in the T-maze reflects circadian rytms. It reached approximately 80 % during the dark phase and dropped down to 20% during the light one (Fig. 3A). We then estimated, for the first 100 trials, the mean probability of choosing i) the left arm in S1, ii) sucrose in S2 and iii) water in S3. We found that mice preferentially chose the most rewarded side, i.e. sucrose for S2 and water for S3 (Fig. 3B). In S1, mice randomly opted for the two arms (i.e. 50% each) at the population level. The evolution of the probability to choose the best option after a bottle swap (Fig. 3B, green or blue curve) suggest a classical reinforcement learning process for tracking the best-rewarded side by trial-and-error. In addition, at the population level, mice showed a decreased return time after choosing the less-rewarded side (Supplementary Fig. 4A) and used a win-stay strategy: they chose the same side after finding the best-rewarded side with high probability, but virtually chose randomly (i.e. around 50%) after missing it (Fig. 3C). A closer examination at the level of individual behaviors revealed that some mice did not alternate in S2 and thus failed to allocate their choices according to the location of the highest reward (Fig. 3D). To identify differences in behaviors, individual choice sequences were thus characterized by four variables that aimed to differentiate choice strategy. Two of these variables (*α* and *β*) were derived from modeling the choice sequence using a classical “softmax” model of reinforcement learning/decision-making. The other two (switch rate noted SW, and slope a) were directly estimated from the choice sequence (see methods). Principal component and clustering analysis distinguished three groups of mice (Fig. 3E): i) G1 mice, characterized by a low switch and virtually no alternation, which always visited the same arm independently of the reward location; ii) G2 mice, which are characterized by an intermediate behavior; and iii) G3 mice, which consistently switched to track higher rewards. The low (LS), intermediate (IS) and high switch (HS) rates of the animals were found to be good indicators for distinguishing the three groups (Fig. 3F). Although the behavior of LS mice may appear suboptimal, this population emerged in most experiments (mean ± sem = 22.1% ± 7.5, n=19/86 mice from 9 experiments).

**Fig. 3:**
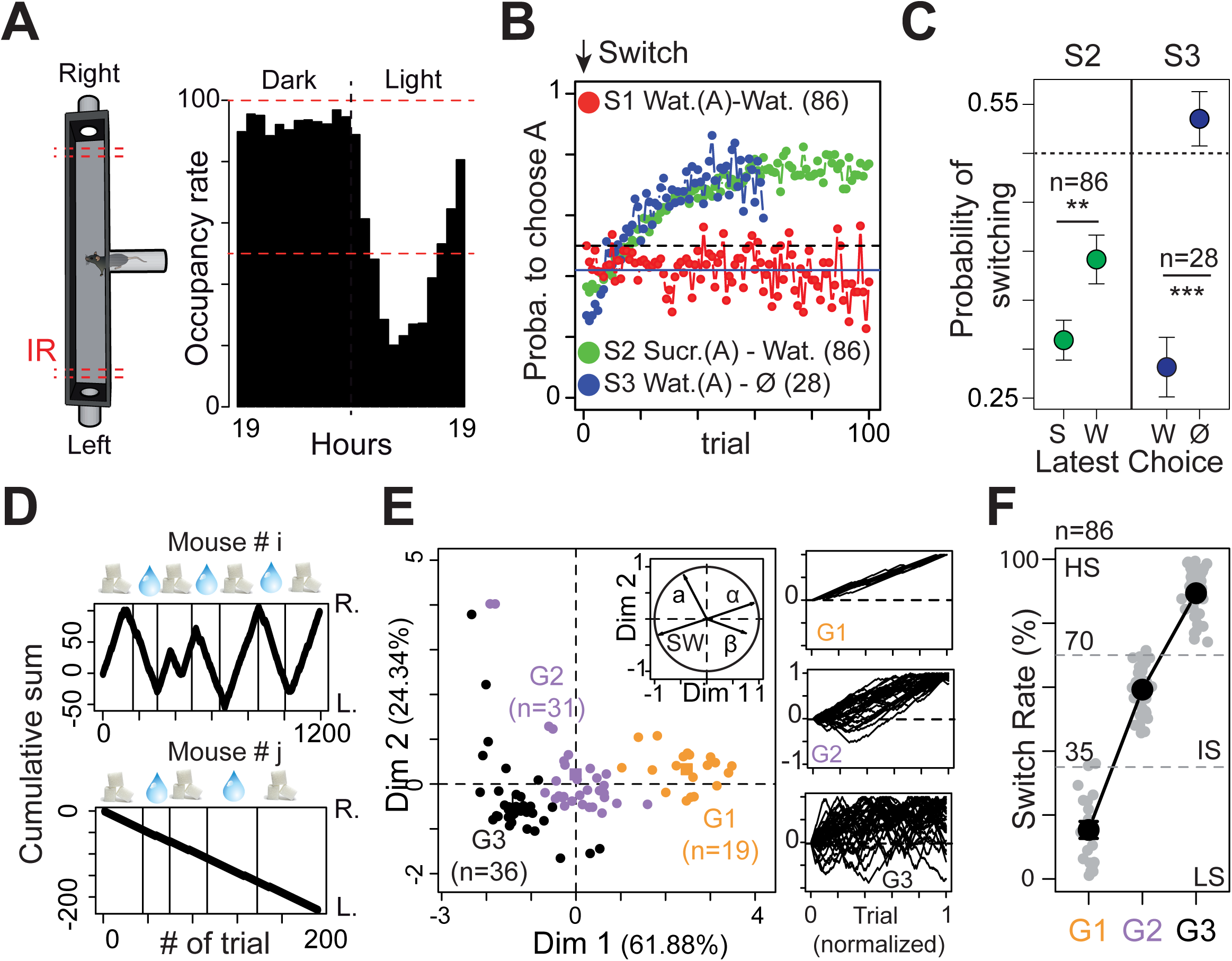
Decision making. (**A**) T-maze occupancy (in %) on a 24h cycle. (**B**) Probability to choose the highest rewarded arm (A) in sessions S1, S2 and S3. For S2 and S3, the first choice corresponds to the one after the bottles have been swapped. (**C**) Win-Stay strategy: probability to switch side when the latest choice (in x-axis) is sucrose (S) or water (W) for S2 and water (W) or nothing (N) for S3. (W=4393 and 675.5, p=0.0012 and p<0.001). (**D**) Cumulative left (L.) or right (R.) turns for two different mice (* i and j), upon water and sucrose bottle swapping in S2 (symbols on top, indicating bottle content on the R. side). (**E**) Principal component analysis based on a, SW; *α* and *β*, (n=86 from 9 experiments) from which we clustered three different groups (G1, G2 and G3). Insets on the right show normalized plots equivalent to (D). (**F**) The three groups are well characterized by their difference in SW (i.e. low (LS), intermediate (IS) and high switch (HS) rates). Data (C, F) are presented as mean±sem; ***p<0.001, **p<0.01, *p<0.05.

### The activity of dopaminergic neurons correlated with individual profiles

After having revealed the existence of various profiles in Souris City, we next aimed at linking cognitive performances in the T-maze with individual traits derived from spontaneous behaviors and with individual neurophysiological activities. Five groups of 10 mice were analyzed. We found that the SW obtained in S2 (Fig. 4A) correlated with other traits of social and non-social spontaneous behaviors (Fig. 4B). Notably, LS mice visited the test zone less frequently than the other groups (Fig. 4B Left) but spent more time in the food compartment (Fig. 4B Middle) or with groups of three or more congeners (Fig. 4B Right). We then assessed whether these phenotypical differences correlated with physiological alterations of specific neural networks, and more specifically the mesolimbic DA system, which is often considered as an important player in personality neuroscience (DeYoung, 2013). Variations in DA have indeed been observed across behavioral traits (Marinelli and McCutcheon, 2014). Moreover, this pathway was shown to encode the rewarding properties of goal-directed behaviors, including social interaction (Gunaydin et al., 2014), and to be a key system in stress-related disorders and addiction (Robison and Nestler, 2011; Russo et al., 2012). Importantly, repeated social defeats produces strong and long-lasting changes within the mesolimbic DA pathway, leading to social withdrawal of defeated individuals (Barik et al., 2013; Berton et al., 2006). To address differences in the DA system between animals, we systematically recorded the activity of DA neurons following an experiment in Souris City. Mice were anesthetized and ventral tegmental area (VTA) DA cell activity was recorded using glass electrodes. DA cell firing was analysed with respect to the average firing rate and the percentage of spikes within bursts (see methods for burst quantification (Eddine et al., 2015; Faure et al., 2014; Ungless and Grace, 2012)). We first compared VTA DA cell activity in mice living in Souris City and in conspecifics living in a standard cage (StC). Both the firing frequency and the bursting activity of VTA DA cells were significantly lower in Souris City compared to StC (Fig. 4C left, Supplementary Fig. 5A-C). Furthermore, when analyzing separately the three groups of mice (LS, IS, HS), an inverted correlation between SW and both the frequency and bursting activity of VTA DA cells was observed (Fig. 4C middle and right, Supplementary Fig. 5D). These results demonstrate a biological inscription, at the level of the midbrain DA system, of the stable and distinctive patterns of behavioral activity that emerged in this complex environment.

**Fig. 4:**
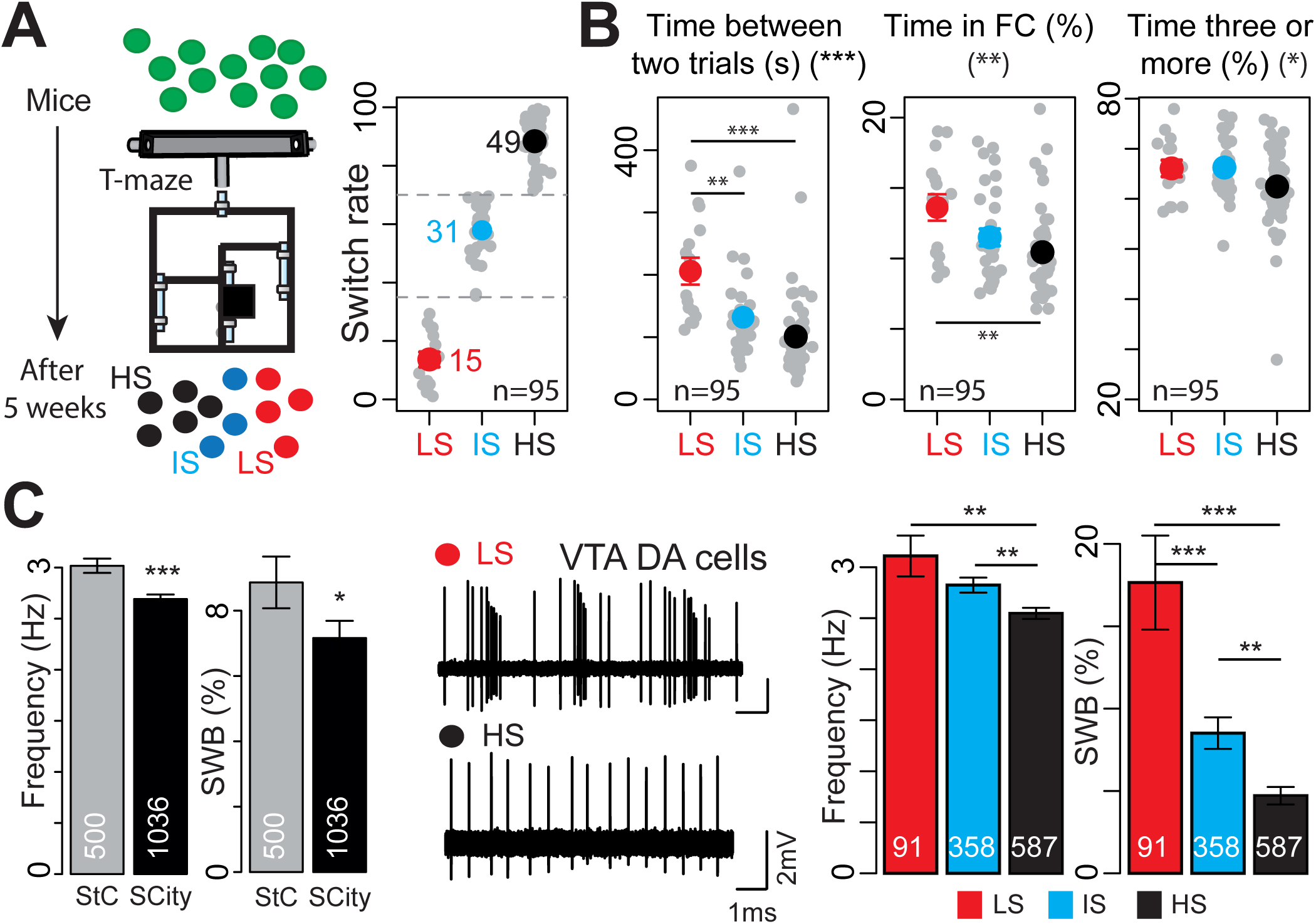
Correlation between specific cognitive behaviors and electrophysiological properties of the DA system. (**A**) Different groups of mice were tracked for 5 weeks in Souris City and classified according to their SW in three groups (i.e. low (LS), intermediate (IS) and high switch (HS) rates). (**B**) Correlation of typical behaviors with SW (from 9 experiments). (Left) Time between two trials in the T-maze (F(2, 92)=12.35, p<0.001; and Tukey post-hoc test). (Middle) Proportion of time spent in FC (F(2, 92)=5.827, p=0.0041, and Tukey post-hoc test). (Right) Proportion of time spent with three or more conspecifics (F(2, 92)=3.177, p=0.04). (**C**) (Left) Spontaneous DA cell activity in standard cages (StC) or in Souris City (SCity) (Frequency and %SWB: W=291920, p<0.001 and W=278010 p=0.015). (Middle) Representative electrophysiological recordings of DA cells from LS (above) and HS mice (below). (Right) VTA DA neuron firing activity of the three groups (F(2, 1033)=8.667 for frequency, F(2, 1033)=26 for %SWB; and Tukey post-hoc test). All data are presented as mean±sem, ***p<0.001, **p<0.01, *p<0.05.

### Social relationships determined both individual profiles and dopaminergic activity

An important question remained, as whether these patterns were irreversible, i.e. related to intrinsic accumulated differences or, conversely, rapidly reversible. We addressed this issue by modifying the composition of two different groups of mice studied in parallel in two Souris City environments (Fig. 5A). During the sucrose versus water session, we used the median SW value to split mice from each Souris City in two populations: the lowest and highest switchers (Step 1). We then mixed the two populations and grouped the lowest switchers from the two environments together, and the highest switchers together. After three weeks of sucrose versus water, we re-evaluated the switching pattern for each mouse (Step 2). Interestingly, distinct switching profiles “re-emerged” within each of the two populations (HS, IS and LS), with no significant difference in the overall distribution of SW before and after mixing (Fig. 5B). Individually, mice that had been relocated (referred to as incomers) to an unknown Souris City decreased their SW (e.g. mouse number #5 in Fig. 5C) whereas mice that did not move (referred to as residents) increased their SW (e.g. mouse number #6 in Fig. 5C). Variation of switching (i.e. SW_step2_ - SW_step1_) was higher in incomers than in residents (Fig. 5D). SW in step 1 was not predictive of SW in step 2: SW of the lowest switchers was homogenous in step 1 (Fig. 5E left) but greatly diverged in step 2, with a clear SW difference between residents and incomers (Fig. 5E right). Finally, we asked whether adaptation of SW was associated with a modification of VTA DA cell firing activity. DA neurons of incomers showed both higher firing rate and bursting activity than those of residents (Fig. 5F, Supplementary Fig. 6A). Altogether, these results suggest that the distinctive patterns of behavioral activity that emerged in this environment are rapidly reversible, and that social relationships can indeed shape behavior and affect the decision-making system.

**Fig. 5:**
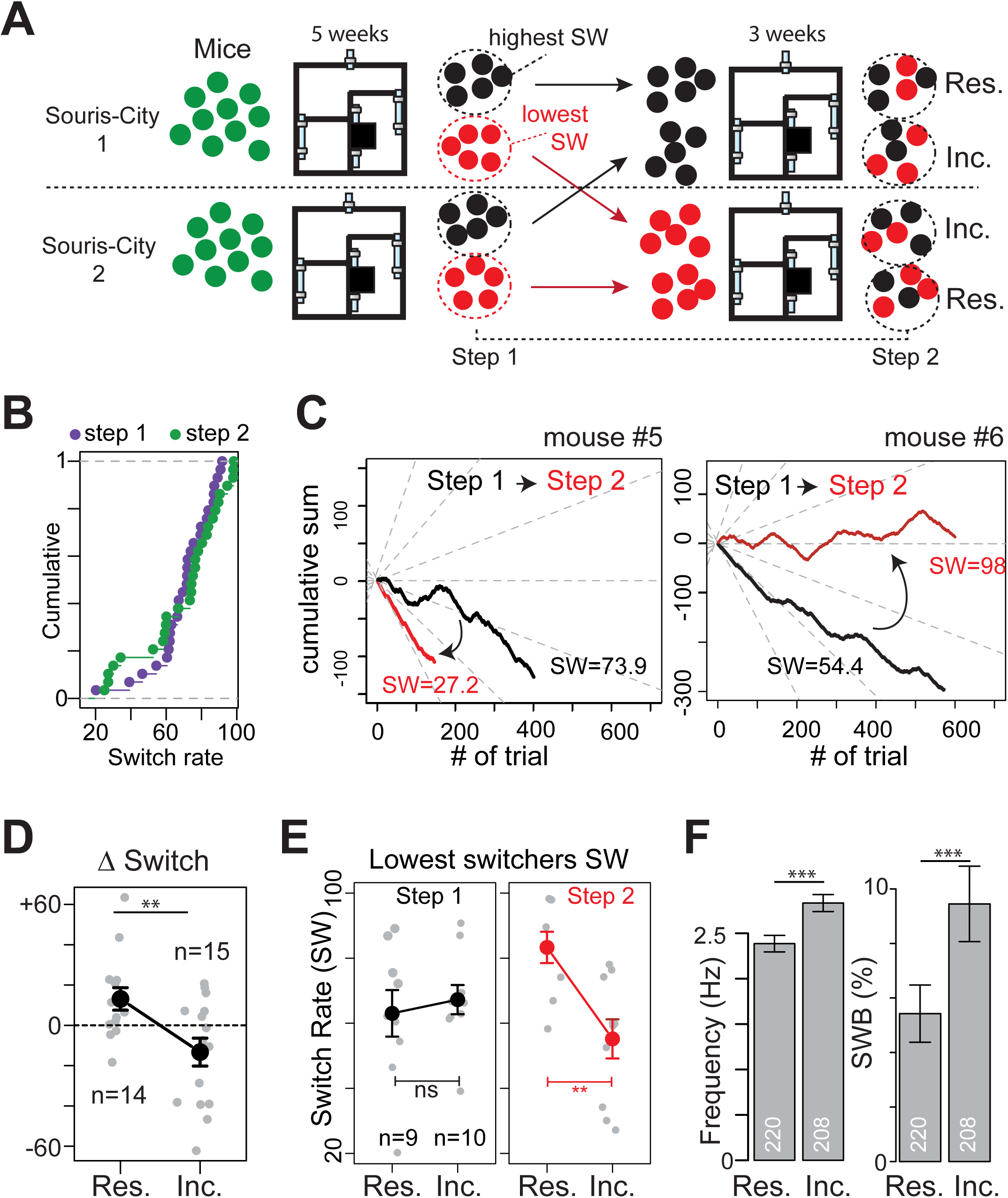
Influence of the group on individual behaviors and on the DA system. (**A**)Experimental paradigm: Two different groups of mice were studied in parallel in two Souris City environments (1 and 2). After five weeks, their switching pattern were evaluated (step 1). Mice from each Souris City were split in lowest (red) and highest (black) switchers. The two populations were then mixed and the lowest switchers from the two environments were grouped together (same for the highest switchers). After three weeks of sucrose versus water, the switching patterns were reevaluated (step 2) for both residents (Res.) and incomers (Inc.). (**B**) Cumulative distribution of SW for steps 1 (purple) and 2 (green) (D=0.2069, p=0.57). (**C**) Cumulative left or right turns for two different mice upon water and sucrose bottle swapping in step1 (Black) and step2 (Red). The incomer mouse #5 switched less, whereas the resident mouse #6 switched more in step2 compared to step1. (**D**) Switch variation between step 1 and 2 μSW) for incomers and residents (two-sample t-test, t(27)=2.9401). (**E**) (left) No difference in SW in step1 between lowest switchers of the two different Souris City, whether they will be subsequently considered as incomers or residents. (Right) SW is different for the same two groups after step2 (Two sample t-test (t=3.5914, **p<0.01)) (**F**) Firing activity of VTA DA neurons from incomers and residents (Frequency and SWB: two-sample Wilcoxon, W=17750 and W=18319 respectively, p<0.001). ***p<0.001, **p<0.01, *p<0.05.

## Discussion

### Large environments and individuation

Groups of mice have complex social structures (Crowcroft, 1966). Social interactions markedly influence a number of behaviors (Larrieu et al., 2017; Lathe, 2004), yet how they affect the development of inter individual variabilities have been rarely addressed in standardized tests. Numerous studies emphasize the needs of using large social housing environments, with automatic testing (Castelhano-Carlos et al., 2014; Sandi, 2008; Schaefer and Claridge-Chang, 2012; Tecott and Nestler, 2004; Vyssotski et al., 2002). Such environments have up until now been mainly used to evaluate strain differences (Endo et al., 2011; Krackow et al., 2010) or test the effect of specific perturbations such as stress on subgroups (Castelhano-Carlos et al., 2014). An essential benefit of automation is that it challenges the classical paradigms consisting in the analysis of average behaviors in distinct groups of animals observed on a short time scale, and puts forward the statistical analysis of individuals recorded in an ecological situation over long time scales. In a relatively stable context, genetically identical animals adjust their behavior over time and situations, yet only within a given range, defining individuality. The notion of individuality thus challenges the idea that behavior of an individual is plastic and thus able to adapt optimally to its environment (Bach, 2009; Bergmüller and Taborsky, 2010; Duckworth, 2010; Sih et al., 2004). For instance, the fact that in our setting two individuals could be classified as either high or low switchers necessarily implies consistency in their decision-making system, and may reflect a limitation to their respective range of adaptation. Our results suggest that this limitation is, on the one hand, strongly linked to local social rules, as evidenced by the experiment where we swapped social environments and, on the other hand, not influenced by local and immediate dynamic of social interactions, since the decision to switch is made in isolation from the congeners.

### Social determinism

Initial variations on a small scale (developmental, epigenetic or micro-environmental) have been proposed to support phenotypic variations on a large scale (Freund et al., 2013; Stern et al., 2017). These small variations are believed to get amplified, resulting in a time-divergence of individual profiles, perhaps due to self-reinforcing effects of past experiences. In this framework, personality emerges slowly and gradually, from small-scale initial individual variations to generate unique phenotypical trajectories. These assumptions do not necessarily imply that personality remains unchanged throughout life (Caspi et al., 2005; MacDonald et al., 2006). Our data shed new light on the role of social behaviors as a factor of divergence contributing to a reorganization of behavior. Social relationships are likely able to amplify initial differences, but can also, as revealed here, trigger rapid and important reshaping of the animal personality and of its DA system, through the dynamic effects of interactions between individuals. These results are compatible with the concept of social niches, which offers an adaptive explanation of the emergence of individuality based on specialization (Bergmüller and Taborsky, 2010). Yet, they also support the idea of a key “social determinism”, in which individuation is decisively determined by social processes and originates from the restriction of the animal capacities to a specific repertoire.

### Specific role of Dopamine

Variations in neuromodulatory functions, including those in the catecholamine and cholinergic systems, might contribute to the process of individuation (MacDonald et al., 2006; Stern et al., 2017). The DA produced in the VTA plays a role in a wide range of behaviors, from processing rewards and aversion to attention, motivation and motor control. The mesolimbic projections participate also in the modulation of social behaviors, as illustrated by genetics studies in human and physiological approaches in rodents (Gunaydin et al., 2014). In the course of a social interaction, an animal must be able to rapidly choose the appropriate behavior, for approaching or avoiding a conspecific. Previous studies demonstrated that the DAergic system undergoes activity-dependent changes (Hyman et al., 2006) that are triggered by “events” occurring during the lifespan of an individual (Faure et al., 2014; Marinelli and McCutcheon, 2014) and that affect basal activity in the long term. The modifications of DA cell activity observed in Souris City may reflect consequences of “social events”. Indeed, it has been shown that the regulation of the DAergic transmission is sensitive to social-stress exposure (Ambroggi et al., 2009; Barik et al., 2013; Cao et al., 2010; Friedman et al., 2014; Morel et al., 2017). Alteration of DAergic activity has also been linked to many motor, motivational or cognitive dysfunctions. In particular, alteration of DA levels has been associated with variations in personality traits and, in the case of tonic DA, with exploration/exploitation trade-of or uncertainty seeking (Frank et al., 2009; Naudé et al., 2016). Furthermore, acutely manipulating VTA DA cell activity using optogenetics (Chaudhury et al., 2013) or pharmacology (Barik et al., 2013), in the context of repeated severe social stress, is sufficient to reverse social-induced stress avoidance. All these results suggest a causal relationship between variations of VTA DA cell activity and the expression of specific behaviors.

### Individuality and susceptibility to psychiatric disease

Finally, our results open new perspectives for preclinical studies on rodent models. Preclinical models usually display high inter-individual variability, but do not focus on “individuals”. For instance, repeated social defeat in genetically identical mice leads to the appearance of depressive-like behavior only in a fraction of susceptible animals, but not in resilients (Krishnan et al., 2007; Russo et al., 2012). Our results indicate that social relationships modify behaviors and circuits in a way that mimics the effects of certain mutations or drugs. The “Souris City” setup thus represents a unique opportunity to address causal relationships between cognitive performance in paradigms relevant for psychiatry and individual traits. Understanding how the social rules amplify the difference in behavioral spectrum displayed by otherwise identical animals will undoubtedly help unraveling the factors influencing the susceptibility of particular populations to psychiatric disorders.

## Materials and Methods

### Animals

8 week-old male C57BL/6J mice were obtained from Charles Rivers Laboratories, France. All procedures were performed in agreement with the recommendations for animal experiments issued by the European Commission directives 219/1990 and 220/1990 and approved by the Comité d’Ethique En Expérimentation Animale n°26. All mice were implanted under anesthesia (isoflurane 3% – Iso-Vet, Piramal, UK), with an RFID chip subcutaneously inserted in the back.

### Souris City setup

#### Setup

“Souris City” combines a large environment (the social cage) where groups of male mice live for extended periods of time in semi-natural conditions, and a test-zone where mice have a controlled access to specific areas for drinking. Souris City was house-designed and built by TSE Systems (Germany). Mice were tagged with RFID chips, allowing automatic detection and controlled access to the different areas. Animals were living under a 12h/12h dark-light cycle (lights on at 7am) and had access to food ad libitum.

The social cage is divided into four compartments: NC, which contains a nest, FC where mice have free and uncontrolled access to food, CC and St to get access to the gate (Fig. 1A, Supplementary Fig. 1). NC, FC and CC are located in a 1m × 1m square, on which St is connected by a tube. These different compartments are equipped with RFID antennas on the floor and are connected through tubes that are equally equipped with antennas. Therefore, each transition from one compartment to the other was associated with a detection of the animal by an antenna.

The social cage is connected to the test zone by a gate, which is a key element of the setup (Fig. 1A). The gate (TSE Systems, Germany) is composed of three doors with independent automatic control (Fig. Supplementary 1B), allowing to select animals and control their access to the test zone. Individuals thus performed the test alone (isolated from their congeners) and by themselves, *i.e.* whenever they wished to and without any intervention from the experimenter. The test consists in a T-maze choice task (Dember and Fowler, 1958). Since the T-maze was the only source of water, animals were motivated to perform the test. The T-maze gives access to two home-cages, one on each side (left and right), with a drinking bottle in each. The bottles contained either water, sucrose or were empty. The system was configured in such a way that animals performed a dynamic foraging task. The reward value of the bottle content could be changed, to evaluate whether mice were able to track the highest reward. Such automation of the task, by minimizing handling and the presence of the experimenter, prevents most limitations of human assessment (i.e. cost and time) and eliminates the risks of stress or disturbance of the animal natural cycle (Castelhano-Carlos et al., 2014; Sandi, 2008; Schaefer and Claridge-Chang, 2012; Spruijt and DeVisser, 2006). Simple rules were used to automatize the test. When a mouse accessed one feeder, the infrared light beam was cut off in that arm, which triggered closing of the feeder on the other side (a Plexiglas cylinder drops in and prevents access to the bottle). Mice had to exit the T-maze to trigger re-opening of the feeders and hence to resume a new trial (Supplementary Fig. 1D). Bottles (for example sucrose- or water-containing) were swapped every 3-4 days.

#### Event detection and storage

Four different kinds of sensors provided automatic data registration in Souris City: RFID antennas surrounding the tubes that connect compartments together (n=14), the gate (n=1), infra-red beam sensors in the T-maze (n=4, 2 on each side) and RFID antennas on the floor (n=16). The IntelliMaze software (TSE Systems, Germany) ran the first three sensors, while TraffiCage (TSE Systems, Germany) controled the floor RFID antennas. These two software programs worked completely independently. IntelliMaze registered a table (.txt file) for each sensor, where each line corresponds to a detection event with the information on animal identity (RFID tag), detection time (millisecond precision), antenna number for the tubes and animal direction for the gate. The TraffiCage software registered detection events as a raw file (.txt file) with the information of animal identity, detection time and antenna number. All these detection events were stored in a database (MySQL relational database hosted by an Apache server), together with spatial and temporal annotation allowing to track the position and activity for each mouse (i.e. mouse number, date, time, antenna number). A web interface coded in php imported the data from the files into the database, linked all the events to the appropriate mouse and created gate sessions. All these events constitute the basic data used for further analysis (see data analysis). R scripts (RMySQL package) were used to extract data from the database.

#### Data Processing

Detection events were used to build various indices and estimators of the animal behavior. The position of the animal was used to calculate its overall activity: i) the proportion of time spent in each compartment, ii) the density of transitions between compartments computed on 24 hours, binned by 10 min periods to evaluate the circadian rhythm, iii) the number of detections for each antenna, and iv) the entropy of each animal. Entropy was calculated from the proportion of time *p* spent in each compartment *i*:

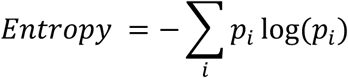

The localization of a mouse relative to others was used to assess the social relationships between mice, e.g. the proportion of time spent alone, with one conspecific or more. We also used detections from both tubes and floor antennas to quantify “chasing episodes” between two mice. Chasing episodes were defined by concomitant (i.e. within a 5s window) detections of the same two mice on at least two consecutive antennas. Antennas were considered consecutive if the first mouse from a concomitant detection on one antenna was detected within a 30s window on another antenna (see Fig. 1.D for schematics). Cumulative curves (entropy and time spent in FC) over sessions represent data from dark phase section (from 7pm to 7am the following day) summed with data from the dark section of the previous days.

#### The T-maze choice quantification

Individual choice sequences (i.e. left or right, Fig. 3) were characterized using four parameters: the switch rate (SW, see above), the slope of the left-right choice (a value close to 1 indicating no switching), the exploratory parameter (*β*) and the learning rate parameter (*α*). We calculated SW for each animal as follows:

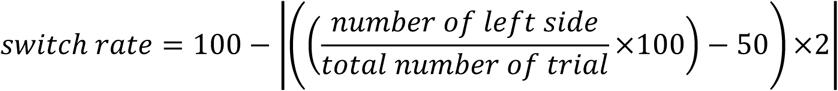

A SW of 100% indicates that the mouse equally chose both sides, while a SW of 0% means that the mouse never switched and always chose the same side. Exploration/exploitation parameters were calculated by fitting the sequence of choices with a standard Reinforcement-Learning/Decision-making model. We used a classical softmax decision-making model where choices depend on the difference between the expected rewards of the two alternatives. This model formalizes the fact that the larger the difference in rewards is, the higher the probability to select the best option will be. Sensitivity to reward difference was formalized by the free parameter *β*. Expectation of reward was adapted through classical reinforcement-learning algorithm, i.e. trial and error, by comparison between the current estimate of action; with *R* (water) = 1, *R* (sucrose) = 2, *R* (nothing) = 0. The value *V_i_* of each action *i* was updated by *V_i_*(*t* + 1) = *V_i_*(*t*) + *αR(t)*, where the free parameter *α* formalizes the learning speed. The softmax choice rule was:

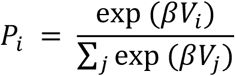

where *β* is an inverse temperature parameter reflecting the choice sensitivity to the difference between decision variables: high *β* corresponds to mice that often choose what they estimate the highest-value arm, while low *β* corresponds to random choice. The free parameters *α* and *β* were optimized using the log-likelihood of the model, on a choice-by-choice basis.

#### Behavioral experiments

The system consists in two parallel and identical setups (Fig.1A, Supplementary Fig. 1) enabling the analysis of up to 10 mice in each of them. In this study, 15 experiments were performed, 12 of which were paired, i.e. executed in parallel in two independent setups. Two setups were coupled (at the St level) for a single experiment, which allowed the tracking of 18 mice. This experiment was used to illustrate some typical results on a larger group of mice (Fig. 2A-C). Overall, 141 mice were tested in Souris City.

### *In vivo* electrophysiological recordings

Mice were anesthetized with an intraperitoneal injection of chloral hydrate (8%), 400 mg/kg, supplemented as required to maintain optimal anesthesia throughout the experiment, and positioned in a stereotaxic frame (David Kopf). Body temperature was kept at 37°C by means of a thermostatically controlled heating blanket. All animals had a catheter inserted into their saphenous vein for *i.v.* administrations of drugs. Recordings were performed using classical technics commonly used in the laboratory (Eddine et al., 2015; Morel et al., 2014). Briefly, recording electrodes were pulled with a Narishige electrode puller from borosilicate glass capillaries (Harvard Apparatus). The tips were broken under a microscope. These electrodes had tip diameters of 1-2 mm and impedances of 20-50 MΩ. A reference electrode was placed into the subcutaneous tissue. When a single unit was well isolated, the unit activity digitized at 12.5 kHz was stored in the Spike2 program (Cambridge Electronic Design, UK). The electrophysiological characteristics of VTA DA neurons were analyzed in the active cells encountered by systematically passing the microelectrode in a stereotaxically defined block of brain tissue including the VTA. Its margins ranged from 3 to 3.8 mm posterior to Bregma, 0.25 to 0.8 mm mediolateral with respect to Bregma, and 4.0 to 4.8 mm ventral to the cortical surface according to the coordinates of Paxinos and Franklin (Paxinos and Franklin, 2004). Sampling was initiated on the right side, and then on the left side. After a baseline recording of 10-15 minutes, the electrode was moved to find another cell. Extracellular identification of DA neurons was based on their location as well as on a set of unique electrophysiological properties that characterize these cells in vivo: (i) a typical triphasic action potential with a marked negative deflection; (ii) a characteristic long duration (>2.0 Ms); (iii) an action potential width from start to negative through > 1.1 Ms; (iv) a slow firing rate (<10 Hz and >1 Hz) with an irregular single spiking pattern and occasional short, slow bursting activity. These electrophysiological properties distinguish DA from non-DA neurons (Ungless and Grace, 2012)

### DA cell firing analysis

DA cell firing was analysed with respect to the average firing rate and the percentage of spikes within bursts (%SWB, number of spikes within burst divided by total number of spikes). Bursts were identified as discrete events consisting of a sequence of spikes such that their onset is defined by two consecutive spikes within an interval <80 ms and they terminated with an interval >160 ms (Eddine et al., 2015; Faure et al., 2014; Ungless and Grace, 2012).

### Statistics

Data are presented as means ± SEM with corresponding dot plots overlaid, as cumulative distribution function, or as boxplot. Data from electrophysiological recording (Fig. 4) are presented as barplot (mean±sem) without dot plots, and their cumulative distributions are presented in supplementary figures (Supplementary Fig. 5 and 6). Statistics for behavioral experiments were carried out using R, a language and environment for statistical computing (2005, http://www.r-project.org). We used a one-way repeated-measures ANOVA followed by a t-test with Bonferroni correction for post hoc analysis to compare the time spent in each compartment through several sessions (Fig. 2C). Consistency over two sessions was estimated by Spearman correlation coefficient (Rho) between several measurements (e.g. proportion of time spent in the compartments) determined in session S1 and S2 (Fig. 2 D, E, F, G). Probability of switching were evaluated using repeated trials (i.e. consecutive entries with a maximum of 20 seconds apart) and were compared using two-sample Wilcoxon test (Fig. 3C). We performed a clustering (bclust function from e1071 package) and a Principal Component Analysis (PCA function from FactoMine package) to define three groups of mice from the T-maze scores (Fig. 3E). We used a one-way ANOVA followed by a Tukey test for post hoc analysis to compare the firing rate and the percentage of DA neuron spikes from LS, IS and HS mice (Fig. 4B). The firing rate and %SWB of DA neurons were compared using two-sample t.test or two-sample wilcoxon test (Fig. 4C Left) or one Way Anova followed by Tukey post-hoc test (Fig. 4C right). SW distribution in mice population were compared using two-sample Kolmogorov-Smirnov test (Fig. 4E left). We calculated the difference between the SW before and after mixing the mice and we compared the incomers with the residents with a t-test or a Wilcoxon test depending on the distribution normality (Fig. 4E right)). The firing rate and %SWB of DA neuron were compared between these two groups with a Wilcoxon test (Fig. 4F).

## Acknowledgements

We thank J. Hazan, E. Ey and F. Tronche for critical reading of the manuscript. This work was supported by the Centre National de la Recherche Scientifique CNRS UMR 8246, the Foundation for Medical Research (FRM, Equipe FRM DEQ2013326488 to P.F), the Bettencourt Schueller Foundation (Coup d’Elan 2012 to P.F.), the Ile de France region (Dim Cerveau et pensée to P.F.), the French National Cancer Institute Grant TABAC-16-022 (to P.F.) and The LabEx Bio-Psy. P.F. and J.M. laboratories are part of the École des Neurosciences de Paris Ile-de-France RTRA network. P.F. and J.M. are members of LabEx Bio-Psy and P.F. is member of DHU Pepsy.

## Author Contributions

N.T. and P.F. designed the study. N.T. and P.F. analyzed the behavioral data. F.M., S.T. and C.N. performed the electrophysiological recordings. C.C., V.O., S.J., L.L.G and S.D. contributed to behavioral experiments. J.N. contributed to analyze the behavioral data. N.D. contributed to create a database. J.M. contributed to the design and the realization of the project. N.T., P.F. and A.M. wrote the manuscript.

## Author Information

The authors declare no competing financial interests. Correspondence should be addressed to.

